# Lung viral infection modelling in a bioengineered whole-organ

**DOI:** 10.1101/2023.02.01.526441

**Authors:** Fabio Tommasini, Thomas Benoist, Soichi Shibuya, Maximillian N.J. Woodall, Eleonora Naldi, Jessica C. Orr, Giovanni Giuseppe Giobbe, Elizabeth F. Maughan, Robert E. Hynds, Asllan Gjinovci, J. Ciaran Hutchinson, Owen J. Arthurs, Sam M. Janes, Nicola Elvassore, Claire M. Smith, Federica Michielin, Alessandro Filippo Pellegata, Paolo De Coppi

## Abstract

Lung infections are one of the leading causes of death worldwide, and this situation has been exacerbated by the emergence of COVID-19. Pre-clinical modelling of viral infections has relied on cell cultures that lack 3D structure and the context of lung extracellular matrices. Here, we propose a bioreactor-based, whole-organ lung model of viral infection. The bioreactor takes advantage of an automated system to achieve efficient decellularization of a whole rat lung, and recellularization of the scaffold using primary human bronchial cells. Automatization allowed for the dynamic culture of airway epithelial cells in a breathing-mimicking setup that led to an even distribution of lung epithelial cells throughout the distal regions. In the sealed bioreactor system, we demonstrate proof-of-concept for viral infection with the engineered lung by infecting primary human airway epithelial cells. Moreover, to assess the possibility of drug screening in this model, we demonstrate the efficacy of the broad-spectrum antiviral Remdesivir. This whole-organ scale lung infection model represents a step towards modelling viral infection of human cells in a 3D context, providing a powerful tool to investigate the mechanisms of the early stages of pathogenic infections and the development of effective treatment strategies for respiratory diseases.

## Introduction

According to the Global Burden of Diseases, Injuries, and Risk Factors Study in 2019, there were approximately 17.2 billion cases of upper respiratory tract infection annually[1]. Since then, the coronavirus disease 2019 (COVID-19) pandemic, caused by severe acute respiratory syndrome coronavirus 2 (SARS-CoV-2), has resulted in an unprecedented challenge to our healthcare system. During the past few years it has become increasingly evident how the development of new drugs is limited by poorly representative cell culture models, and inter-species differences in the pathogenesis of infectious diseases. While testing drugs on monolayer cell cultures is the traditional way of assessing the efficacy of antiviral treatments, such methods lack organ specific structure including 3-dimensional (3D) cell interactions with the lung extracellular matrix (ECM) and may not be ideal to examine the pharmacokinetics in the physiological condition[2]. As such, the development of an infection model of a human engineered respiratory tissue that recapitulates the 3D *in vivo* environment may improve our ability to model early viral infection of epithelial cells.

When considering the respiratory system, attempts at reproducing the complexity of lung microenvironment *in vitro* have included organoid modelling. For example, we and others have used organoid systems to reproduce *in vitro* some specific aspects of COVID-19, such as the GI symptoms[3][4]. However, organoids are still limited by missing lung-specific ECM interactions and architecture and fail to represent more complex tissues and organs. Viral infections have also been investigated at the microscale in airway-on-a-chip devices. Huh et al. have created a lung microfluidic chip to reproduce a functional alveolar-capillary interface of the human lung[5]. The device mimics physiological organ-level functions, including pathogen-induced inflammatory pathways and responses to cytokine exposure. Breathing-type movements affect acute pulmonary cell toxicity and proinflammatory activity as cyclic mechanical strain accentuates toxic and inflammatory responses of the lung to nanoparticles. Similarly, F. Zhang et al. managed to obtain a reliable reconstruction of tissue layers including alveolar epithelium and microvascular endothelium in a microfluidic system[6]. The synergistic effects of the epithelial and endothelial interfaces on the stimuli resistance verified the importance of creating complex tissue microenvironments *in vitro* to explore pollution-involved human pathology. Moreover, L. Si et al. modelled human airways on a chip through a two-channel microfluidic cell culture device composed of an air channel and a blood channel[7]. The air channel is lined by highly differentiated human primary airway cells cultured under an air-liquid interface, while the blood channel is lined by human pulmonary endothelial cells. Influenza A was introduced to the air channel to mimic the airborne route of transmission, and drugs and antibodies could be perfused via the blood channel to mimic drug treatments.

While microfluidic devices do not incorporate complex stoichiometric cell matrix interactions, this can be overcome through the use of organ-scale bioreactors. To date, the most promising method to produce a whole organ scaffold is decellularization, which consists in the removal of the cellular component from an organ while preserving the extracellular matrix (ECM)[8]. Whole organ perfusion decellularization of the lung has been previously described[9]. In particular, these scaffolds feature a preserved micro- and macro-structure, as well as the ECM components and the different vascular, alveolar and proximal/distal airway compartments. Several detergent-based and other physical/chemical decellularization protocols have been described for murine[10][11][12] and large animal models[9][13][14][15][16][17][18]. These showed different degrees of decellularization efficiency, matrix preservation and cellular material removal. Moreover, the decellularization process features an intrinsic variability due to the donor variations such as age, associated conditions and organ preservation. Further variation can occur during the process of recellularization, leading to low data reproducibility among different laboratories. Similarly, different dynamic culture strategies have been described for lung recellularization including diffusion[19], dynamic rotating wall vessels[20], airway ventilation[12], or both airway ventilation and vascular perfusion[10][11]. Among these, airway ventilation provided the most promising results, indeed it mimics what occurs *in vivo*, making this approach the first choice for a dynamic system.

In order to enhance reproducibility of the engineered organ, an automated process that could standardise both decellularization and recellularization processes would be beneficial. Beside the repeatability of the process, automation can reduce contamination and was shown to result in more consistent decellularized matrix quality, in terms of residual DNA, as well as collagen, elastin, laminin and glycosaminoglycan levels[18]. Automation will also facilitate clinical translation, where GMP guidelines require traceable and reproducible processes, and will offer a uniquely safe closed system to investigate airway infections and test innovative therapies. Few reports have investigated automation of the perfusion decellularization process, however, with most focused on specific steps such as the monitoring of the flow rate or pressure[21], or the selection of the decellularization reagents[18].

Here, we describe an innovative system for the automatic decellularization and recellularization of a whole-organ lung scaffold. The automated system delivers the decellularization and recellularization of rat lungs, providing airway ventilation and perfusion. Dynamic recellularization leads to a uniform single-layer cellular distribution throughout the organ, similar to native tissue, while maintaining the phenotype of airway basal cells. Moreover, we demonstrate the infection of the bioengineered lung with respiratory syncytial virus (RSV) and the efficacy of the broad-spectrum antiviral Remdesivir in this model.

## Results

### Bioreactor design and operation

Lung decellularization, recellularization, infection and sample collection were all performed thanks to an automatic, closed and compact bioreactor system (Figure 1A, Supplementary figure 1A). The setup is composed of an electrical compartment and a hydraulic circuit. The fluid distribution system has two different designs for decellularization and recellularization (Figure 1B). The decellularization setup allows lung perfusion with various chemical reagents by inflation and deflation through the trachea, using a series of six electrovalves and two peristaltic pumps. The recellularization setup allows lung inflation and deflation with culture medium, as well as recirculation of medium that the lung is immersed in, using a separate “reservoir” bottle, one electrovalve and one peristaltic pump per lung. Flow rates and durations of fluid movement to and from the lung were chosen so as to gently inflate the lungs close to their maximum capacity (Figure 1C,D, Supplementary figure 1B), so that fluids could reach all parts of the tissue. As a reference, we considered that for rats the total lung capacity is around 11.3 ml, whereas the vital capacity is 8.4 ml[23]. The electrical compartment of the bioreactor is the core of its automation. The Arduino board, which allows programming of the circuit through the Arduino programming language, was connected to a series of stacked Adafruit motor shield boards. These motor shields, which allow control of higher voltage motors as used in our setup, were in turn connected to the main components of the hydraulic circuit. The code was first written in the Arduino integrated development environment (IDE) and then uploaded to the Arduino board, which initiated hydraulic circuit automation. Different codes were used for lung decellularization and recellularization (Supplementary figure 1C,D). In order to improve operational efficiency, two identical bioreactors were built. This also allowed us to run several experiments in parallel. Leakage and sterility tests were performed to guarantee hydraulic seal preservation in the circuit and complete sterility for the entire duration of the experiment (data not shown).

**Figure 1.**
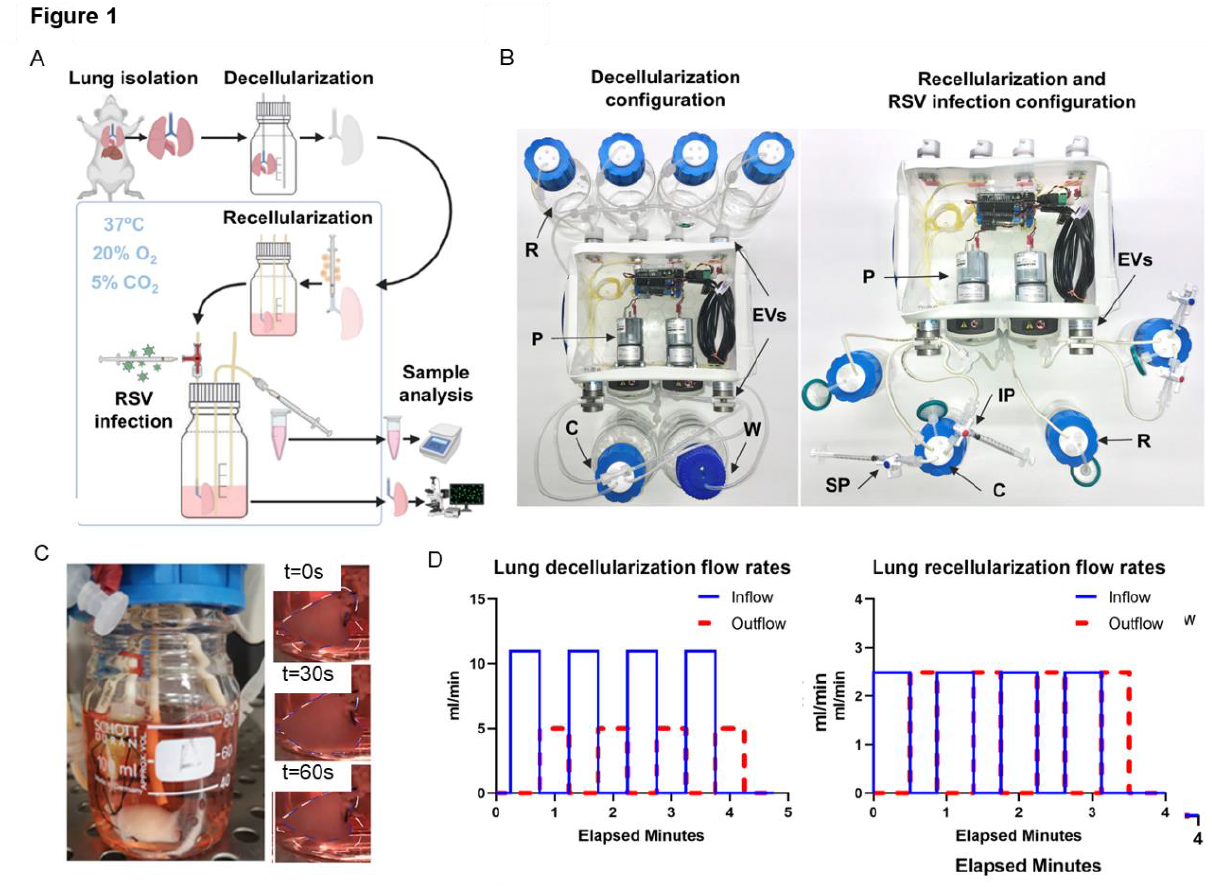
Whole lung engineering system. **A**. Schematic of the experimental process: lungs are harvested and then decellularized using the decellularization configuration of the system then the lung is recellularized using the recellularization configuration which is also used to administer RSV infection via dedicated ports. **B**. Detailed configuration of the (left) decellularization configuration (left) and the recellularization and RSV infection configuration (right). In the decellularization configuration, four reservoirs are connected to the system, while decellularization reagents are pumped in the chamber using pumps (P) and electrovalves (EVs) then discarded into the waste container (W). The recellularization and infection configuration allows the culture of two lungs housed in two separate chambers (C). Media is stored in two sterile reservoirs (R). RSV infection and following analysis is performed via injection ports (IP) and sampling ports (SP) respectively. **C**. Image of bioreactor’s lung chamber during the dynamic culture. Time-lapse of lung breathing during a whole ventilation cycle (total duration of 60 seconds), showing the volume changings of the organ between inflation (white line) and deflation (blue line). **D**. Representative graphs of the lung inlet (red) and lung outlet (pink) flow rates during dynamic breath decellularization (left graph) and during dynamic culture (right graph).

Freshly harvested rat lungs were cannulated through the trachea and decellularized through a detergent and enzyme treatment (DET) protocol[9] using the automated bioreactor system (Figure 1A). This protocol originally included distilled H_2_O, Sodium Deoxycholate (SDC) and DNAse solution. However, during several tests, an interaction between SDC and DNAse solutions was observed by the appearance of a white viscous solution when these solutions were mixed. This seemed to be dependent on the SDC concentration and temperature; in fact, it has been observed that this phenomenon occurred at 4°C, if SDC concentration was around 0.125% and 0.0625%, whereas no gelation occurred at room temperature (data not shown). Hence the SDC concentration as a decellularization reagent was decreased to 2% and a set of washes of the scaffold with PBS during decellularization was implemented to minimise the contact between SDC and DNAse. Furthermore, the entire procedure was performed at room temperature. This optimised decellularization protocol was then automatised. The decellularization setup (Figure 1B, left, Supplementary figure 1C) is composed by four selecting valves (EVs on the back of the bioreactor) connected to decellularization reagents reservoirs (R), two pumps(P) to drive the decellularization fluids in and out the cannulated lung and two valves (EVs on the front of the bioreactor) to control the inflow and outflow of the fluids from and to the organ and the bioreactor chamber (C). The “open”/ “closed” status of the reagent selection valves is related to the solution required for the specific step of the protocol. When the selection valve is open, the corresponding fluid is pumped towards the lung bottle and so on for the other fluids. The decellularization reagents were instilled through the rat trachea by mimicking breaths (inspiration/expiration), so the scaffold is gently inflated and deflated (Figure 1D). At the same time, after each decellularization cycle, the pumps are activated for 1 minute together, in order to wash out the accumulated reagents from all lines and replace it with the new reagent, according to the protocol, before the beginning of the next inflation.

The recellularization system (Figure 1B, right, Supplementary figure 1D) differs slightly from the decellularization one. This system consists of two medium-filled bottles (reservoir (R) and lung chamber (C), each filled to 40ml) linked by airtight silicone tubing. The forward or backward rotation of a peristaltic pump (P) onto the tubing allows either the perfusion or retrieval of medium from the engineered lung. In order to avoid lung airway collapse and damage due to pump-induced negative pressure, more volume is perfused into the lung than it is retrieved. Therefore, to ensure medium level control, the switching of a selecting valve (EVs) allows the peristaltic pump to empty any excess medium in the lung bottle. The tube inlet used to that end is placed at the desired level of the medium. If the medium level rises above this level, the fluid is removed from the lung chamber and pumped to the reservoir. Conversely, if the level of the medium is equal or slightly under air only will be aspirated, leaving the medium level unchanged. Medium levels were chosen so that lung lobes were covered but the trachea was left out of the medium. This conformation allowed the trachea to remain straight thanks to the weight of the lung, so as to avoid tracheal kinking and subsequent blockages. In addition, a support made of autoclavable plastic was designed and added to the hydraulic circuit to keep the lung from the medium-control tube and avoid its aspiration. The infection setup (Figure 1B, right) is very similar to the recellularization one. The main changes involve the adding of an infection port (IP) and a sampling port (SP). The former was added to allow safe inoculation of the virus within the lung, while the latter was added to allow daily sampling of spent medium to measure and analyse viral particles (Supplementary Figure 1A). Specifically, for the infection port a needle-free valve was used to prevent any leakages at the moment of virus inoculation through the trachea. This valve seals the syringe completely, thanks to a rubber spring part located inside the main structure of the valve, avoiding any spillages. This is a critical point when the operator is dealing with the virus inoculation. For the sampling access, a three-way valve was connected to a tube with access to the lung bottle medium.

### Whole-organ decellularization

The system uses 6 valves and 2 pumps to control the flow of decellularization reagents to and from the trachea of rat lungs (Figure 2A). Scaffolds were analysed after either 4 or 9 decellularization cycles (Figure 2B). After 4 cycles, the macroscopic appearance of the lung was evenly translucent white and airway branches were visible. Further tests were performed at the microscopic level to assert the extent of the lung’s decellularization as well as the preservation of the ECM, comparing native to cycle 4 and cycle 9 decellularized lungs. DAPI and Haematoxylin and Eosin (H&E) staining showed that no visible nuclei could be detected in the decellularized lungs after 4 decellularization cycles (Figure 2C). Then, images from H&E staining and a scanning electron microscope (SEM) showed that, after 4 and 9 decellularization cycles, small alveolar, bronchiolar and putative capillary structures were still preserved at a microscopic level (Figure 2C). This was confirmed with Methylene Blue dye injection in the trachea, which showed no macroscopic leakages throughout the airway tree structure (Figure 2D). In addition, micro-computed tomography (μCT) imaging confirmed that the overall integrity of the airways and alveoli were preserved at a macroscopic level (Figure 2E). Performing 9 decellularization cycles rather than 4 with cyclic inflation/deflation of the lung (Figure 2F) was crucial for DNA removal from the scaffold. Indeed, DNA quantification measured 91136 ± 170 ng/mg nucleic acids on fresh lung tissue, 866 ± 147 ng/mg at cycle 4 and a much lower value of 176 ± 53 ng/mg after 9 cycles (Figure 2G). Parenchyma-to-alveolar space was evaluated resulting in no statistically significant difference among fresh (31 ± 6%), cycle 4 (24 ± 2%) and cycle 9 (23 ± 4%). Similarly, there was no difference in the distribution of the alveolar sizes. (Figure 2H, I). Moreover, mechanical testing showed that Young’s modulus of decellularized lungs was not statistically different from the one of fresh tissue (Figure 2J).

**Figure 2.**
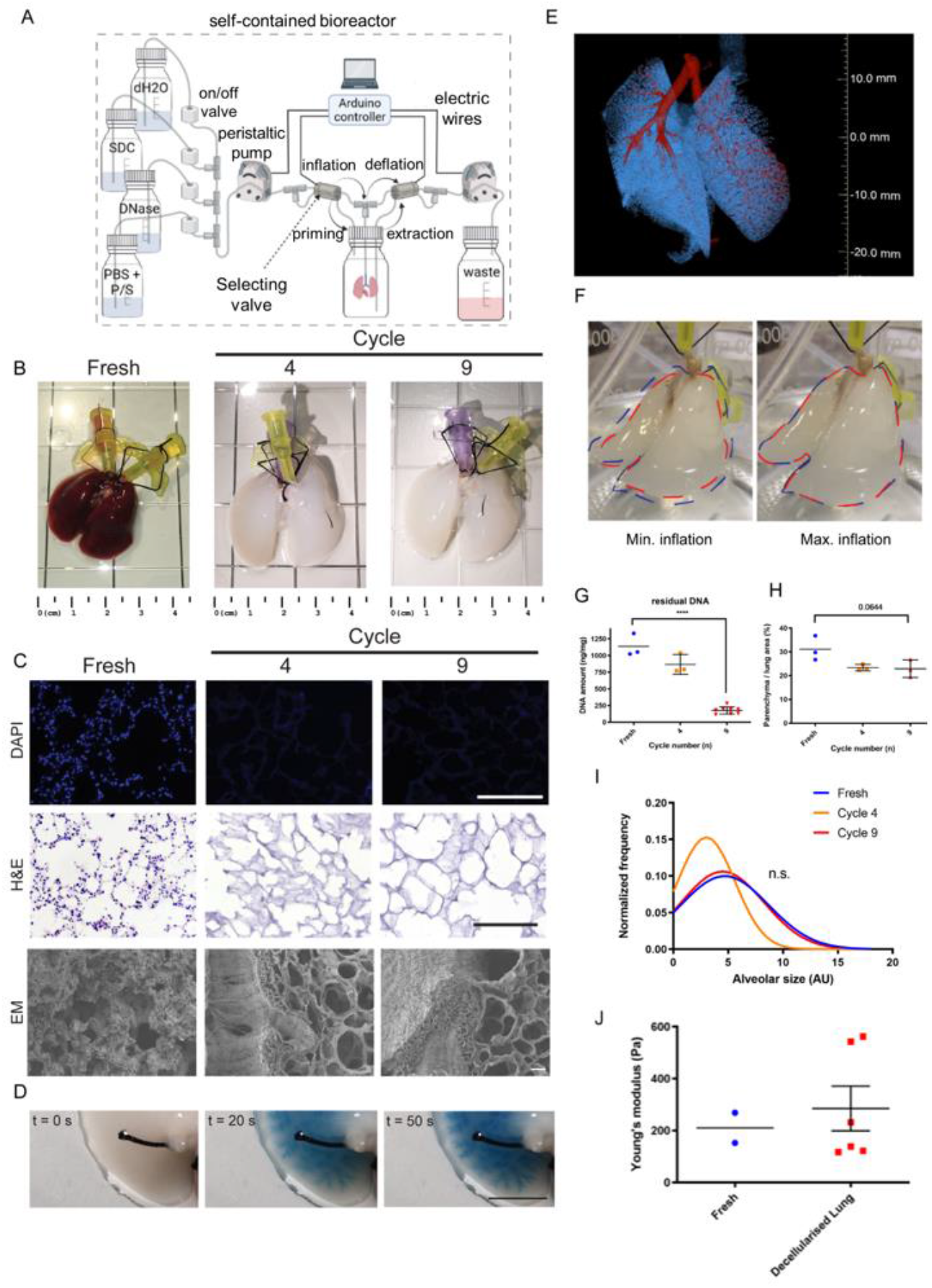
Whole lung decellularization. **A**. Schematic of the bioreactor setup for the whole rat lung decellularization. Decellularization reagents are cyclically perfused intratracheally through the continuous inflation and deflation of the lung controlled automatically by an Arduino controller. **B**. Macroscopic images of the decellularized lung before and after 4 or 9 cycles of decellularization. **C**. Representative images of nuclear staining and H&E assay of the decellularized lung before and after 4 or 9 cycles of decellularization (top and middle line) show efficient cell removal after decellularization as early as Cycle 4. Electron Microscopy analysis (bottom line) confirms preservation of tissue structure upon decellularization. Scale bar 200μm. **D**. Perfusion with dye shows preservation of airways up to the distal areas of the lung. Scale bar 200μm. **E**. Micro-CT scan analysis reveals the preservation of the airways and vasculature after decellularization. **F**. Representative images of the decellularized lung during minimum inflation (red line) and maximum inflation (blue line) show effective expansion of the lung during the dynamic process. **G**. DNA quantification of the decellularized before and after 4 or 9 cycles of decellularization show significant DNA removal at Cycle 9. One way ANOVA; ***p<0.005. **H**. Quantification of histological staining reveals that the percentage of parenchyma to lung area is comparable before and after decellularization. One way ANOVA. **I**. Comparable alveolar size distribution have been observed between fresh and decellularized tissue. **J**. Young’s modulus measurement shows preservation of mechanical stiffness after decellularization.

### Characterisation of decellularized lung ECM composition

We subsequently investigated the composition of the decellularized lung extracellular matrix (ECM), comparing it to fresh tissue. The collagen fraction relative to overall weight was found to increase from fresh tissue (1 ± 0.1μg / mg wet tissue) to cycle 4 (5 ± 1μg / mg wet tissue) and cycle 9 (12 ± 3μg / mg wet tissue) (Figure 3A). Collagen preservation was also confirmed through multiple staining. Picro-sirius red (PR) and immunofluorescence for collagen I and collagen IV highlighted maintenance of both types of collagen lining the bronchi, bronchioli, airway interstitium and alveoli (Figure 3B,C). Interestingly, plane polarized light showed yellow birefringence in structures such as vasculature, and large bronchiole, rich in large collagen fibres such as collagen type I. Likewise green birefringence was observed in the alveolar parenchyma as well as large bronchiole and vasculature which are rich in thin/reticular collagen fibres such as collagen type III respectively (Figure 3C). The maintenance of these optical properties from fresh tissue to decellularized scaffolds is a further proof of the maintenance of collagen integrity. However, sulphated glycosaminoglycans (sGAGs) were less retained during decellularization. sGAGs content in the fresh tissue (0.139 ± 0.003 μg/mg) showed a significant 3-fold reduction between the fresh tissue and cycle 4 (0.057 ± 0.003 μg/mg), although there was no statistical significance between cycle 4 and cycle 9 (0.048 ± 0.008 μg/mg) (Figure 3D). Alcian Blue (AB) staining confirmed partial preservation of sGAGs across the alveolar and vascular parenchyma (Figure 3E). In addition, elastin content underwent a very significant (p<0.001) 3-fold reduction from the fresh tissue (31.7 ± 2.4 ng/mg) to cycle 4 of decellularization (12.3 ± 3.2 ng/mg). Similarly, a modest but significant (p<0.05) reduction was observed between the elastin content of the acellular tissue between cycle 4 and cycle 9 (7.7 ± 1.6 ng/mg) (Figure 3F). Despite the reduction, elastic fibres maintained their distribution, lining the alveolar, bronchial and vascular walls as observed by Elastic Van Gieson (EVG) staining (Figure 3G). Masson Trichrome (MT) and MOVAT staining further highlighted the preservation of collagen fibres within decellularized ECM, as well as confirmed efficient cell removal (Figure 3H). Finally, laminin was shown to be well preserved in the alveolar, bronchial and vascular parenchyma of decellularized lungs (Figure 3I).

**Figure 3.**
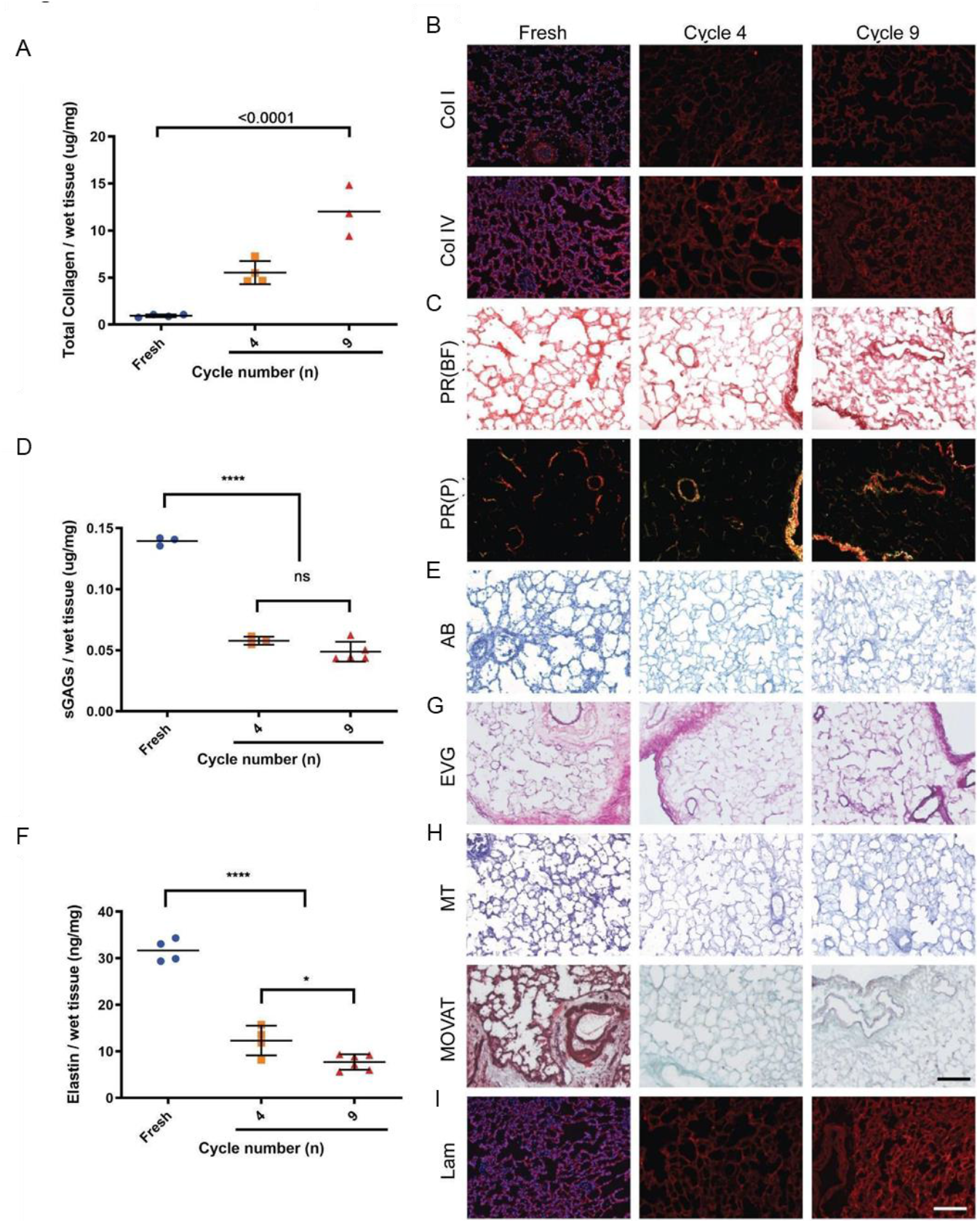
Decellularized lung composition characterization. **A**. Collagen to tissue weight ratio is increased, mostly due to cell removal. **B**. Collagen I and Collagen IV staining shows preservation of both collagen types and preservation of the overall structure. **C**. GAGs quantification shows a reduction after decellularization. **D**. Alcian blue staining shows preservation of GAGs distribution. **E**. Elastin quantification shows a reduction after decellularization. **F**. Elastica Van Gieson staining shows that elastin structure throughout the lung is preserved. **G**. Laminin IF staining shows preservation after decellularization.

In addition, the presence and amount of residual decellularization detergent within decellularized scaffolds was assessed, as it might negatively affect seeded cell survival. This was assessed by measuring the amount of sodium deoxycholate leached into decellularized lung PBS washes, using spectrophotometry. The first PBS wash following SDC treatment was able to decrease the SDC concentration in the collected effluent by 10 fold (0.2%). Then, the second set of PBS washes after the DNAse phase was able to decrease it further by 10 fold (0.02%) (Supplementary Figure 1E).

### Whole-organ recellularization

Decellularized whole lungs were manually seeded with a minimum of 20 millions of either A549 cells or HBECs through trachea. Then, they were left in static culture from overnight to 18 hours for cells to adhere before being put under dynamic culture. Dynamic culture was performed using the recellularization bioreactor system (Figure 4A) described before, which drives pump-powered inhalations and exhalations through the lung’s cannulated trachea. Static culture conditions were also performed as controls.

**Figure 4.**
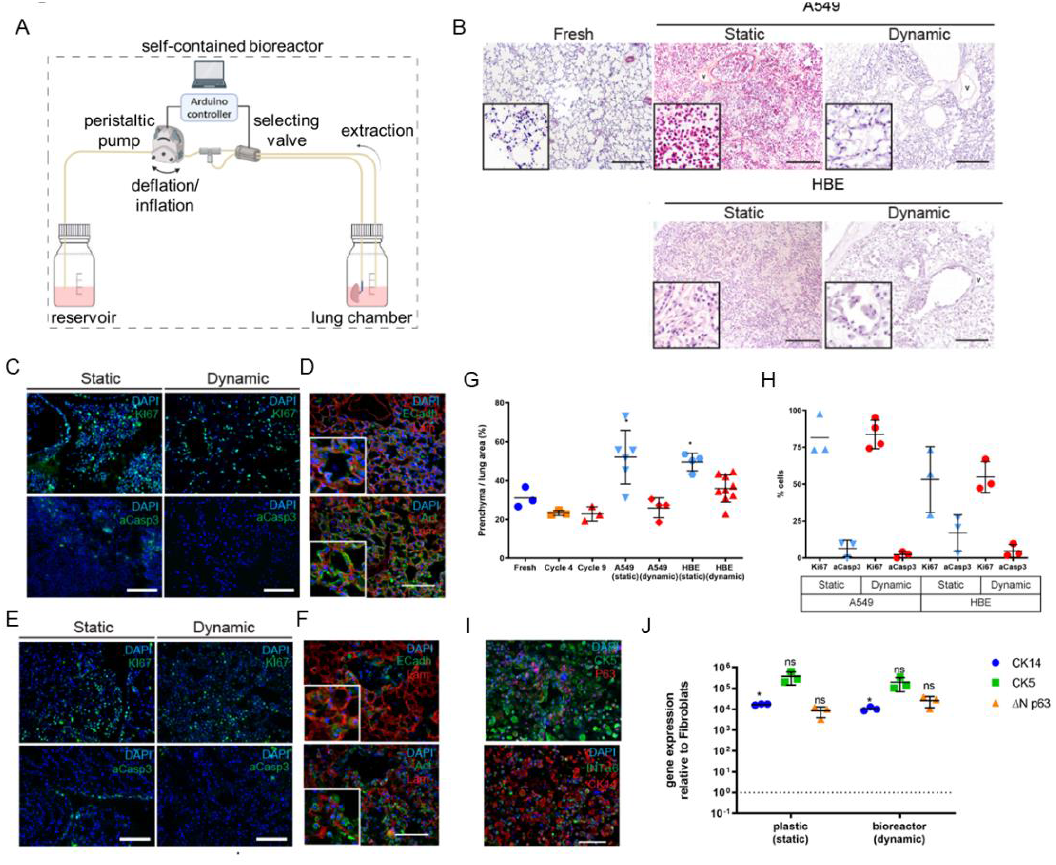
Whole lung recellularization. **A**. Schematic of the bioreactor setup for the whole rat lung recellularization. **B**. H&E staining after recellularization with A549 alveolar cells and Human Bronchiolar epithelial cells (HBECs) cultured under static or dynamic conditions, compared to fresh tissue. Widespread cell distribution, similar to native tissue, can be observed for both cell types. **C**. Immunofluorescence analysis of A549 cells for proliferation and apoptotic markers Ki67 and aCasp3, respectively, show high proliferation and reduced apoptosis in all conditions. **D**. Dynamic recellularization with A549 cells show a proper cellular organisation and expression of E-Cadherin. **E**. Immunofluorescence analysis of HBECs for proliferation and apoptotic markers Ki67 and aCasp3, respectively, show high proliferation and reduced apoptosis in all conditions. **F**. Dynamic recellularization with HBECs show a proper cellular organisation and expression of E-Cadherin. **G**. Dynamic recellularization results in a parenchyma over lung area ratio comparable to native tissue and lower in respect to static recellularization. **H**. Quantification of the percentage of cells positive for KI67 and Caspase3 in the different condition. **I**. HBEC dynamic recellularization shows expression of lung epithelial markers. **J**. Gene expression analysis of selected lung epithelial genes. CK14: Cytokeratin 14; CK5: Cytokeratin 5; ΔN p63: DeltaN p63.

After 5 days under static culture conditions, lung ECM demonstrated a clear overcrowding with both A549 cells and HBECs, even so that the alveolar structures could not be recognised. Conversely, dynamic culture conditions showed an organised cellular distribution along the ECM and a patent airway space, similar to the native control lung tissue structure (Figure 4B). The percentage of parenchyma over lung area was evaluated as a measure of the cumulative space occupied by the cells and scaffold. Cell proliferation and viability was not negatively affected by dynamic culture conditions. Assessment of proliferating A549 cells, using Ki67 staining, showed rates of 81 ± 14% in static versus 83 ± 10% in dynamic conditions. For HBECs, Ki67 staining was 53 ± 22% for statics versus 54 ± 10% for dynamic conditions. Viability of A549 cells, assessed with aCasp3 staining, was quantified at 6 ± 5% in static and 2 ± 2% in dynamic conditions. For HBECs, aCasp3 staining was 16 ± 12% in static versus 4 ± 4% in dynamic conditions (Figure 4C, E, H). The percentage of parenchyma in the static conditions (A549: 52 ± 13%; HBECs: 49 ± 4%) was significantly higher (p<0.05) than the fresh tissue (31 ± 5%) while the percentage of parenchyma in the dynamic conditions (A549: 25 ± 5%; HBECs: 35 ± 6%) was similar to fresh tissue (31 ± 5%) (Figure 4G).

Under dynamic culture conditions, cells expressed the epithelial marker E-Cadherin in close proximity to the basal lamina. Similarly, cell shape as observed by Actin staining seemed to accommodate the geometry of the alveolar ECM (Figure 4D). HBECs, cultured in dynamic condition showed adhesion to the broncho-alveolar ECM and formed a homogeneous monolayer over the basal lamina (Figure 4F). HBECs cells cultured under dynamic conditions for 5 days in their maintenance medium co-expressed airway basal cell markers Cytokeratin 5 (CK5), Tumor Protein p63 (p63), Cytokeratin 14 (CK14) and Integrin-α6 (INT α6) (Figure 4I). The gene expression levels of the airway basal cell markers CK14, CK5 and ΔNp63 were at similar levels to the HBECs maintained *in vitro* on tissue culture polystyrene (TCPS) as a control (Figure 4J).

### Whole-lung infection

Engineered lungs were then subjected to RSV infection and treatment with the broad-spectrum antiviral Remdesivir, shown to inhibit RSV replication in vitro[23]. An adapted recellularization setup was used for this purpose, designed to facilitate RSV infection, Remdesivir administration and daily medium sampling while protecting the operator and bioengineered lung from cross-contamination (Figure 5A). In order to understand the influence of RSV and Remdesivir on the lungs, four independent experimental conditions were set (Figure 5B). The four experimental arms, which were executed simultaneously, consisted of dynamic culture of an engineered lung with or without RSV infection (RSV+ and RSV-respectively) and with or without Remdesivir treatment (REM+ and REM-respectively). Firstly, decellularized lungs were seeded with 26 million HBECs and cultured dynamically for 4 days to become “engineered” lungs. Then, RSV (or PBS for control arms) was injected into the lung through an injection port and left to incubate at 37°C for 1 hour in static culture, with the lung kept above the medium for that duration. Then, Remdesivir (or DMSO for control arms) was added to the lung’s medium through the sampling port and dynamic culture was resumed. 2D culture controls in standard polystyrene tissue culture wells were also set up, following the four experimental arms.

**Figure 5.**
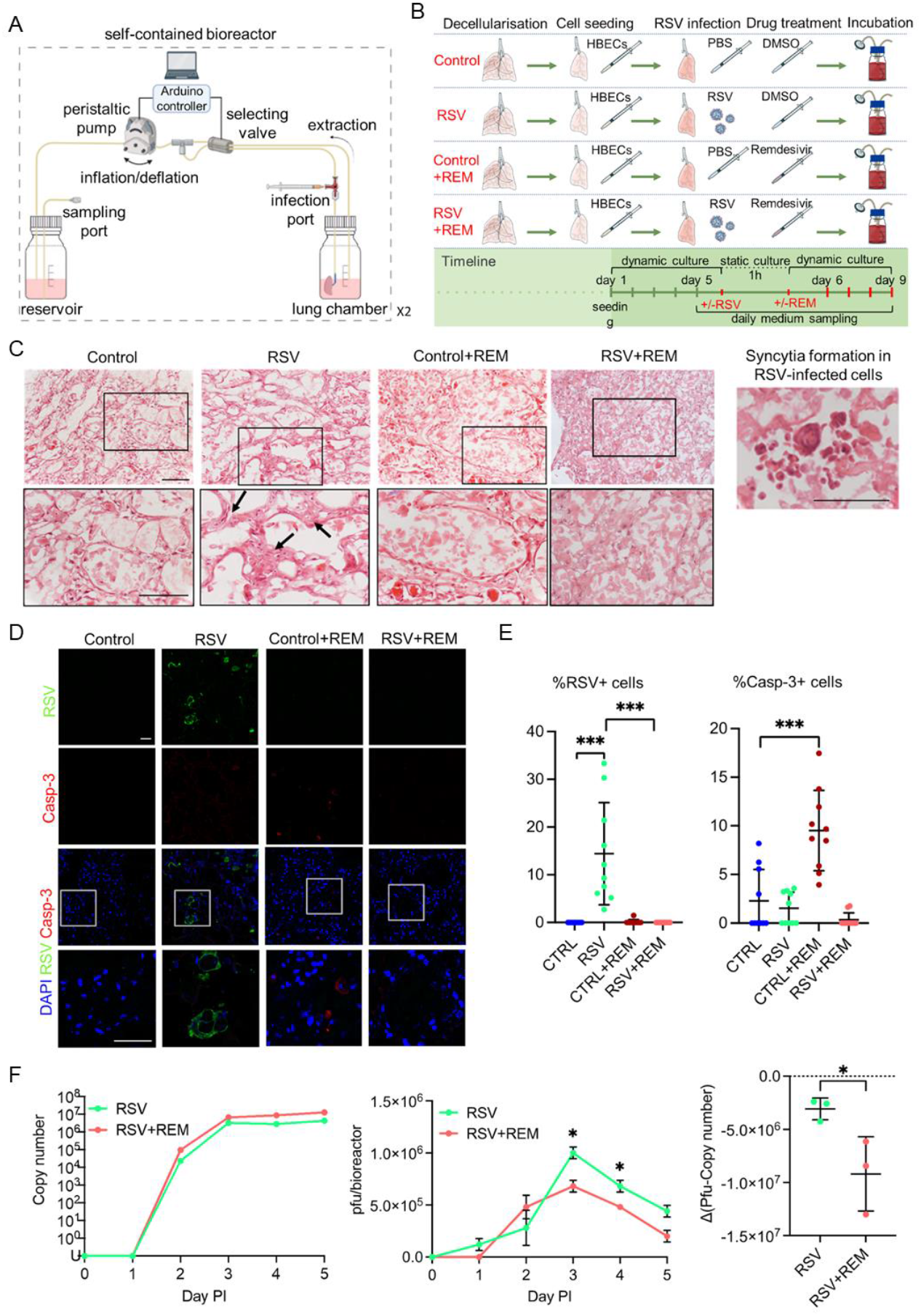
Engineered lung infection. **A**. Schematic of the bioreactor setup for the engineered lung infection and medium sampling. This includes the addition of ports for the safe injection of the viral suspension into the lung and the retrieval of virus-contaminated supernatants. **B**. Experimental setup for the recellularization and infection of the engineered lung with an RSV virus in the presence or absence of the antiviral drug Remdesivir. A non-infection condition and a treatment with Remdesivir without RSV infection have been included as controls. **C**. Haematoxylin and Eosin analysis of the engineered lungs cultured under the four experimental conditions. Black arrows show a thickened parenchyma in the RSV condition that can be observed in the higher magnification pictures (row below). Scale bar 100μm. **D**. Immunofluorescence analysis of RSV and cleaved Caspase-3 proteins in engineered lungs cultured under the four experimental conditions show RSV-positive cells in the infected lung with no Remdesivir and the presence of apoptotic cells (Casp-3). Scale bar 50μm. **E**. Quantification of the immunofluorescence analysis in D shows a significant increase in the percentage of RSV-positive cells in the RSV infection condition (left). The percentage of Caspase 3-positive cells is significantly increased in engineered lungs cultured with Remdesivir only. T-test; ***p<0.005. **F**. Analysis of the viral titer of the supernatants collected from the RSV-infected engineered lungs without or with Remdesivir, at different days post infection (PI). Quantitative RT-PCR of RSV viral load in media samples measured by N protein copy number. Infectious particles measured via Plaque-forming unit (pfu) assay, calculated for all circulating media within a bioreactor (80mls) (pfu/bioreactor) (central). The difference between pfu and copy number (right). Points show mean ± SD, n=3. Multiple t-test; *p<0.05.

Haematoxylin and Eosin stains of engineered lungs cultured under the four experimental conditions (Figure 5C) showed that cells in the untreated RSV-infected arm tended to accumulate and form multi-layer aggregates, pointed by black arrows, while cells were distributed as monolayers in the three other conditions (RSV-REM-, RSV-REM+, RSV+REM+). In a separate experiment, in RSV-infected bioengineered lungs, we were able to observe a large multi-nuclear single cell, which is characteristic of RSV-induced cell syncytia (Figure 5C, right). Immunofluorescence staining of RSV and Caspase 3 were performed on slices from engineered lungs cultured under the four experimental conditions. Consistent positive RSV signals could only be observed in the RSV+REM-condition (Figure 5D, E, Supplementary figure 2D). Then, Caspase 3 signals in the RSV-REM+ condition were seen to be significantly higher (p<0.001) than the RSV-REM-control, suggesting that unbound Remdesivir at the used concentration could be toxic to cells. Prior experiments comparing RSV+ to RSV-engineered lungs showed similar results to the experiment above (Supplementary Figure 5A, B, C), one exception being Casp-3+ cells being significantly higher in the RSV+ than the RSV-arm (p<0.005).

After lung infection and Remdesivir treatment, the medium circulating in the bioreactor was sampled daily. The sampled medium was analysed for viral copy number (specifically N-gene copies) by rt qPCR and for viral infectivity using a plaque forming assay (Figure 5F). Viral copy number of the supernatants collected from the RSV-infected engineered lungs without or with Remdesivir displayed a similar increase post-infection (PI) (left). However, plaque-forming units (pfu) were significantly lower (p<0.005) in the RSV+REM+ condition than the RSV+REM-condition at PI day 3 and 4 (central), also confirmed in the difference of pfu and copy number (right) (p<0.005).

## Discussion

A tissue-engineered whole organ would represent a powerful tool for both disease modelling and regenerative medicine applications. Aside from the possibility of challenging transplantation shortage, a whole engineered organ is a prime tool to model airway disease mechanisms, whether these are related to infection or to test personalised medicine approaches.

Here, we describe a tissue-engineered whole lung system used for modelling viral infection. This model uses decellularized rodent lungs which have been recellularized using human epithelial cells and subsequently infected with RSV. Results show the feasibility of RSV infection and the efficacy of Remdesivir in lowering viral infection. The whole model, from decellularization up to the infection process, is delivered in and by an innovative automatic and dynamic culture system.

The first step of the automated process is a whole-organ decellularization of a rat lung via a detergent-enzymatic protocol. A staple of decellularization is the complete removal of the cellular and nuclear material while preserving the extracellular matrix[8]. The automatized process allowed full decellularization of the organ, which retained its extracellular matrix composition and mechanical properties. As previously described also by our group[9][24][25], we exploited an airway perfusion to deliver the decellularization reagents in the lung, which allows their direct delivery to the proximal and distal regions of the lobe. Several manual lung decellularization protocols have been used for different species ranging from mice to humans. Nevertheless, the variability among decellularization protocols in terms of concentration of detergents, pH as well as the innate differences in the lung matrix composition and structure across species clearly calls for automation and standardisation[26]. To overcome this limitation, here we describe a fully automated setup able to deliver a dynamic decellularization, recellularization and infection of the whole organ. The decellularization configuration allows for an automatic exchange of the decellularization reagents according to a user defined protocol involving up to four reagents, which is the maximum number of reagents commonly found in the literature. Our system-operated decellularization resulted in good preservation of the macro- and micro-structure at both the bronchioalveolar and vascular level. Moreover, the system allows the delivery of the protocol in a closed environment that minimises the risk of contamination and reduces operator-related risks.

Following decellularization, our system could shift into a recellularization setup which has been used to engineer a re-epithelialized lung thanks to a dynamic culture protocol which mimicked physiological breathing. Dynamic culture conditions prompted a uniform single-layered recellularization by HBECs, which was not achieved using static culture. Indeed, dynamic conditions promoted the formation of an epithelial monolayer, which lined the small airways, the terminal bronchiole and the alveoli, leaving patent airspace, similarly to what has been observed in fresh tissue. On the other hand, the decellularized lung cultured under static conditions appeared to be congested.

Strikingly, dynamic culture conditions promoted cell proliferation and lowered apoptosis, while retaining the cellular phenotype. These beneficial effects of mechanical liquid ventilation could be explained by the added fluid flow. This could improve cell viability thanks to a better diffusion of nutrients and the removal of waste material and dead cells within the decellularized lung.

Using an adapted version of the recellularization setup which includes injection and sampling ports, we were able to leverage this whole lung engineered construct to perform a proof-of-concept experiment of viral infection by targeting airways epithelial cells, the primary target for viral infection and replication in the lung[27]. To do so, we delivered an RSV viral solution intratracheally to infect the bronchial cells in a safe and controlled manner through the closed bioreactor system.

Moreover, we tested the antiviral properties of Remdesivir, by administering the drug in the engineered lung upon infection and comparing with controls. Results showed the feasibility of RSV infection and the potential to screen antiviral agents, such as Remdesivir, in the whole-organ model, providing for the first time a 3D lung whole-organ model of viral infection, while overcoming current limitations related to the use of miniaturized systems[2][5][6][7].

This innovative, whole-organ lung model represents a promising tool to investigate the mechanisms of the emergence of pathogenic infections and the development of effective treatment strategies for respiratory diseases in the future. This 3D *in vitro* culture system could serve as a drug-screening platform or as a gene therapy model. Of note, the recent Covid-19 outbreak highlighted the necessity for *in vitro* culture platforms that can speed up and scale out drug discovery and mechanism studies. The 3D organ in-vitro system here presented could represent a powerful tool to address these issues, answering a need that has been clearly pointed out by reviews in the field[28]. Moreover, the automated and dynamic decellularization-recellularization system can be easily exploited on other organs, opening the possibility of creating other whole organ models.

Finally, our model presents some limitations that can be eventually addressed in further studies. First our cellular compartment is limited to epithelial cells, which, despite being the prime target for airway viral infections, are just a part of the whole infection mechanism that also involves the immune system and the vascularisation. It would be extremely interesting to implement these cellular compartments in future studies. Moreover, HBECs correspond to basal cells, which represent just a part of the several phenotypes that compose the airway epithelium; it would be important to include all the other epithelial phenotypes such as ciliated, goblet or club cells. From a fluid dynamic perspective, two remarks could be done. First the RSV, as well as other airway viral infections occur via airborne particles, thus air inflation should be used to mimic this process, while in our system the virus is administered via culture medium. Second, the vasculature should be the way to deliver media that maintain cells alive and eventually mimic the administration of systemic drugs.

In conclusion, despite the above-mentioned limitations, our system represents the first example of a whole-organ, dynamic and automated system to model a lung infection. This system will prove useful in providing insight in the mechanism of viral infection and eventually defining personalised medicine approaches.

## Supporting information

Suppplementary informations

## Acknowledgments

This research was supported by the Longfonds Voorhen Astma Fonds BREATH Consortium (Award 552269 to P.D.C.) and the NIHR Great Ormond Street Hospital Biomedical Research Centre. T.B. is supported by a NIHR Great Ormond Street Hospital BRC Studentship. R.E.H. was a Wellcome Trust Sir Henry Wellcome Fellow (WT209199/Z/17/Z) and is supported by a NIHR Great Ormond Street Hospital BRC Collaborative Catalyst Fellowship. C.M.S and M.W were supported by grants from BBSRC (BB/V006738/1). F.M. is supported by a NIHR Great Ormond Street Hospital BRC Catalyst Fellowship. P.D.C. is supported by National Institute for Health Research (NIHR-RP-2014-04-046). The views expressed are those of the authors and not necessarily those of the NHS, the NIHR or the Department of Health.

## Materials and Methods

### qPCR

Tissue was chopped with a scalpel and transferred into an eppendorfEppendorf tube. TRI Reagent (Cat. # T9424, Sigma Aldrich, UK) was added and samples were vortexed. Samples were then stored in −80°C until needed. When thawed, chloroform (Cat. # 372978, Sigma Aldrich, UK) was added and the samples were centrifuged. The clear phase (supernatant) was collected and one volume of ice cold 70% Ethanol added. The mixture was then loaded on spin columns from the RNAeasy mini kit (Cat. # 74104, Qiagen, UK). Several washes were performed with RW1 and RPE buffer supplied with the kit, according to manufacturer’s recommendations. The RNA was then quantified and the quality checked by measuring the Absorbance ratio Abs_λ=260 nm_ / Abs_λ=280 nm_ and Abs_λ=260 nm_ / Abs_λ=230 nm_. Complementary DNA (cDNA) was generated from 2μg of total RNA per reaction using High Capacity cDNA Reverse Transcription kit (Cat. # 4368814, Thermo Fisher, UK). qPCR was performed using 10ng of template and 200μM of each forward and reverse primer (Supplementary methods) with SYBR green PCR master mix (Cat. #4334973, Applied Biosystem, UK), following the manufacturer’s recommendations. For TaqMan primers (Thermo Fisher, UK), listed in supplementary methods, qPCR master mix was prepared according to manufacturer’s recommendations, using 10ng of template for each replicate. The expression of each gene was measured in triplicate in a 96 well plate PCR plate loaded on a StepOnePlus Real Time PCR system (Applied Biosystem, UK).

### Organ retrieval and preparation

Procedures on animals were performed in accordance with local approvals and under licences released from the UK Home office. Adult Sprague-Dawley rats weighing approximately 250 – 350 g were euthanized by CO_2_ inhalation and exsanguination in accordance with local animal guidelines and approvals (in the UK, Home Office guidelines under the Animals-Scientific Procedures-Act 1986). Median sternotomy was made, and the cervical muscles were divided along the midline to well expose the trachea. The thymus was removed. The superior and inferior vena cava were separated. The main trunk of pulmonary artery was cannulated via the right atrium and secured with a 5/0 silk suture, through which pulmonary circulation was flashed with 2mM/L EDTA. The pulmonary artery was then separated and the branches of the pulmonary vein were carefully divided to free the lungs. The trachea was cannulated with a 24G catheter, which was secured using a 5/0 silk suture, and transected at just above the cricoid cartilage. Cannulation of the trachea was confirmed by injecting 1-3ml of PBS to see the lungs well perfused. The lungs were then extracted and placed on a petri dish containing PBS and Penicillin-Streptomycin (P/S).

### Decellularization protocol

The modified version of a previously published detergent enzymatic treatment (DET) protocol[10] was adopted to decellularise the lungs. The lungs were inflated and deflated (“breath”) through the trachea with enzymes and reagents. The duration of a breathing cycle of inspiration or expiration was 30 seconds. One cycle of decellularisation consisted of 4 breaths with double distilled water, 4 breaths with 2% Sodium Deoxycholate (SDC) (Cat. # D6750, Sigma Aldrich, UK), 8 breaths with PBS supplemented with 1% penicillin/streptomycin solution (PBS - P/S), 4 breaths with 2,000 kU/L DNAse (Cat. # D5025, Sigma Aldrich, UK) in a solution of 9 g/L NaCl, 1 g/L CaCl_2_, and 8 breaths with PBS. Decellularisation was completed by 9 cycles of treatment.

### Decellularization/Recellularization automatic setup

The automatic decellularization setup is composed of an electrical component and a fluid distribution system. Regarding the electrical components, an Arduino UNO R3 (Cat. # A000066, RS, UK) board was programmed through the Arduino IDE coding environment. The board was interfaced with a series of stacked Adafruit Motorshield boards, which deliver current to the different motors of the fluid distribution system (pumps and valves). The fluid distribution system is composed of a series of 2-Ways Normally Closed pinch valves (Cat. # 225P011-21, NResearch, UK) and 3-Ways pinch valves (Cat. # 225P091-21, NResearch, UK), which would accommodate a silicone tubing of ID 1/16 inch (Cat. # TBGM101, NResearch, UK) connected with Luer adapter tees and barbed hoses (Coleparmer, UK). Fluid was moved by the powering of two 114 FDC peristaltic pumps (Cat. # 010.5DP0.00A, Watson-Marlow, UK). For the recellularization, airtight Pharmed tubes (Cat. # R6502-16BPT, Saint Gobin, UK) were used to minimise bubble formation.

### Micro-computerised tomography (micro-CT)

Decellularized lungs were injected with Microfil contrast agent (Flow Tech, Carver, MA, USA) into the pulmonary artery and air was injected into the trachea. The specimen was then submerged in water and scanned using a μCT apparatus. The specimen was secured using foam supports, Parafilm M (Bemis, Oshkosh, WI, USA) and carbon fibre rods to ensure mechanical stability during micro-CT examination. Micro-CT images of the specimen were acquired using an XT H 225 ST microfocus CT scanner (Nikon Metrology, Tring, UK) with the multi-metal target set to Tungsten. X-ray energy and beam current settings were 80 kV and 88 μA respectively. Exposure time was 500 ms, with the number of projections optimized for the size of the specimen (number of pixels covered within area of interest x 1.5) and one X-ray frame per projection. Projection images were reconstructed using modified Feldkamp filtered backprojection algorithms with proprietary software (CTPro3D; Nikon Metrology) and post-processed using region-growing tools in VG Studio MAX 2.2 (Volume Graphics GmbH). Isotropic voxel size was approximately 15 micrometres.

### Anionic detergent SDC quantification

The concentration of Sodium Deoxycholate (SDC) used in the decellularization procedure was determined using an optimised and published protocol[9]. A 2% SDC solution was prepared in PBS - P/S and further diluted in PBS to generate the standard samples. Briefly, 1 or 10μl of the wash effluents and standard samples were mixed with 0.0125% methylene blue (Cat. # 03978, Sigma Aldrich, UK). After vortexing, chloroform was added to the effluents and standard samples at a 1:2 ratio. Effluents and standard samples were then left for 1 hour at room temperature. 150μl of the chloroform phase were extracted and the absorbance was read at 595 nm with a Tecan Infinity microplate spectrophotometer. A standard curve was generated by plotting the mean Absorbance of each standard sample against the SDC concentration of the standards. Linear regression was used to generate a best-fit linearized curve through the points on the graph. The amount of SDC in the wash effluents could then be calculated.

### OCT embedding and slides preparation

Samples were fixed by immersion in 4% PFA overnight at 4°C, washed in PBS post-fixation and left at 4°C for 30 minutes in a solution of 10%, and then of 15% sucrose in distilled water. Samples were then left at 4°C overnight in a solution of 30% sucrose. The following day, samples were embedded in Optimal Cutting Temperature (OCT) tissue embedding compound (Cat. # 62550-01, Electron Microscopy Science, UK) and frozen by immersion of OCT cassette in isopentane (Cat. # 277258, Sigma Aldrich, UK) over liquid nitrogen or over dry ice. Samples were then stored at −80°C. When needed, sample tissues were sliced into 10μm sections using a Leica Cryostat and further stored at −20 up to −80°C.

### Immunofluorescence of OCT embedded cryosections

OCT embedded tissue slices were immersed in PBS for 15 minutes and the tissue was permeabilized using 0.3-0.5% Triton X-100 for 10 minutes. The areas including the tissue sections were encircled with hydrophobic Dako Pen (Cat. # S200230-2, Agilent, UK) to minimise the reagent volumes used. Tissues were blocked using 5% Normal GS in PBS for 30 minutes to 1 hour at room temperature. The tissues were then immersed in solutions containing primary antibodies overnight at 4°C and in solutions containing secondary antibodies at room temperature for one hour. A list of the antibodies used can be found in the supplementary methods. Antibodies were dissolved in 1% Normal Goat Serum in 0.1% Triton X-100.

### Alveolar size quantification

Random fields of H&E stained lung slices were photographed and images were manually processed in Fiji (ImageJ) as follows. Image background was subtracted using a rolling ball algorithm of 20 pixels radius. The images were then filtered using a median filter of 1 pixel. Up to 60 alveoli were visually identified and selected with the tracing tool with 2 pixels threshold. Regions of interest (ROI) were then automatically generated and the areas were computed using a scale bar conversion between real distance and pixels. Results from the quantifications of different samples were compared using Prism 6 software.

### Scanning electron microscopy (SEM)

Native and acellular lung tissue was fixed in 2.5% Glutaraldehyde (Cat. # 808326, Sigma Aldrich, UK) in PBS for at least 18 hours. Samples were processed by Dr. Claire Crowley. Samples were washed in PBS, cut in fragments and immersed in cryoprotectant solution (25% sucrose, 10% glycerol in 0.05M PBS) for two hours. The samples were snap frozen in liquid nitrogen and fractured. Following the thawing of the fragments in the cryoprotectant solution at room temperature, samples were washed in PBS and fixed in 1% OsO_4_ for 2 hours. Fragments were then dehydrated in a graded ethanol-water series to 100% ethanol, dried using critical point CO_2_ and mounted on aluminium stub. Samples were then coated with a thin layer of Au/Pd using a Gatan ion beam coater. Several fields at different magnifications were recorded using a Jeol 7401 FEG scanning electron microscope. Biological replicates of different samples were also assessed.

### Percentage of alveolar parenchyma

To assess the integrity of the macro architecture of the lung parenchyma in native and decellularized lung specimens, images of a minimum of three random fields of H&E staining were acquired. At this point Images were analysed in Fiji. Background was subtracted with a rolling ball of 50-pixel radius, to also increase the contrast between the parenchyma and the alveolar areas. RGB images were then converted to 8-bit and the threshold was automatically set to highlight the tissue, leaving the airways untouched. The parenchyma area was selected and the percentage of the selected tissue was calculated relative to the total area of the field.

### Inspection of decellularized matrix integrity

Decellularized rat lung and isolated lobes were perfused with a solution of 0.1% Methylene blue solution (Cat. # 03978, Sigma Aldrich, UK) in PBS either from the trachea or pulmonary artery, at a constant flow. A video was taken over the course of the injection and still frames were acquired at regular intervals using Adobe Premiere Pro CS6.

### Lung stiffness measurement

Samples of fresh and decellularized rat lung were cut in pieces of about 0.2cm^3^ and secured to the bottom of a 3cm dish using cyanoacrylate glue. The samples were then rinsed two times in PBS and imaged using Piuma Nanoindenter (Optics11, UK) with a spherical tip with a diameter of 50μm. Young’s modulus was computed for at least 3 randomly selected regions of the matrix.

### Total DNA quantification

Tissue samples were isolated from native and decellularized biopsies and dissected into approximately 25mg pieces. Tissue pieces were then finely minced and gDNA was isolated using DNeasy Blood and Tissue kit (Cat. # 69504, QIAGEN, UK) according to the manufacturer’s protocol. Briefly, samples were lysed in proteinase K and a digestion buffer and loaded in a mini spin column. After extensive alcohol washing, the double stranded DNA was eluted and quantified using a NanoDrop spectrophotometer (Nanodrop, US). Absorbance at 230nm, 260nm and 280nm were used to estimate the purity and yield of nucleic acid. For all quantifications, at least three replicates were obtained. All samples were processed in parallel to minimise user-dependent variability between samples. The mean value for total gDNA yield was then computed for each of the fresh and decellularized tissue samples and differences were assessed for statistical significance using Student’s t-test.

### Total Collagen quantification

Approximately 30 mg native and acellular tissue samples were weighed, finely minced and incubated for 18 hours in 6M HCl at 95°C. Hydrolysed samples and standards were processed using the Total Collagen kit (Cat. # QZBtotcol2, QuickZyme Biosciences, The Netherlands). The lysates were then incubated with a chromogenic solution staining hydroxyproline residues and left to develop colour at 60°C for 1 hour. Samples and standards were run in duplicates and absorbance was read using a Tecan Infinity microplate spectrophotometer with band pass filter of 555nm. A standard curve was generated by plotting the mean absorbance of each standard against the collagen content of the standards. Linear regression was used to generate a best-fit linearized curve. The amount of collagen in the original samples could then be computed.

### Sulphated glycosaminoglycans (sGAGs) quantification

Native and acellular tissue samples were weighed (approximately 60 mg), finely minced and digested for three hours in a papain extraction reagent (0.01% in 0.2 M phosphate buffer, pH 6.4) at 65°C for 18 hours, with occasional vortexing. Digested samples and standards were processed using the Blyscan Sulphated Glycosaminoglycan assay (Cat. # B1000, Biocolor, UK) according to the manufacturer’s recommendations. Samples were incubated with 1,9-dimethyl-methylene blue dye and reagents from the assay. Samples and standards were run in duplicates and absorbance was read using a Tecan Infinity microplate spectrophotometer with band pass filter of 595 nm. A standard curve was generated by plotting the mean absorbance of each standard against the collagen content of the standards. Linear regression was used to generate a line of best-fit. The amount of sulphated glycosaminoglycan in the original samples could then be computed.

### Elastin quantification

Native and acellular tissue samples were weighed (approximately 30mg), finely homogenised and complete solubilisation of elastin was performed for two hours in 0.25M oxalic acid at 100°C. Digested samples and standards were processed using the Fastin Elastin Assay (Cat. # F2000, Biocolor, UK), according to the manufacturer’s recommendations. Extracts were incubated with 5,10,15,20-tetraphenyl-21H, 23H-porphine tetrasulfonate dye. Samples and standards were run in duplicates and absorbance was read using a Tecan Infinity microplate spectrophotometer with band pass filter of 520nm. A standard curve was generated by plotting the mean absorbance of each standard against the elastin content of the standards. Linear regression was used to generate a line of best-fit. The amount of elastin in the original samples could then be computed.

### Haematoxylin and Eosin staining

OCT embedded slides were reconstituted in PBS for 15 minutes and rehydrated using serial 5-minute descending baths of ethanol (100%, 90%, 70%). Slides were then washed in running tap water and stained with Hematoxylin (Cat. # H3136, Sigma Aldrich, UK) and then Eosin (Cat. # E4009, Sigma Aldrich, UK) for 5 minutes each. Slides were then dehydrated in 5-minute ethanol baths of ascending percentages (70%, 90%, 100%) followed by Histoclear (Cat. # HS-200, National Diagnostics, USA) before being mounted with DPX mounting medium (Cat. # 06522, Sigma Aldrich, UK).

### Masson Trichrome staining

Tissue sections were stained with Masson Trichrome stain (Cat. # 361670, RAL diagnostic, France) according to the manufacturer’s recommendation. Briefly, slides were rehydrated using ethanol baths of decreasing percentages (100% to 70%) and stained in Mayer’s Haemalum for 10 minutes, rinsed in running water for 4 minutes, followed by staining in Ponceau-Fuchsin solution. The slides were then rinsed with two baths of 1% Acetic acid in water, treated with 1% phosphomolybdic acid and stained in Aniline Blue solution. Sections were then dehydrated in ethanol baths of ascending percentages (70% to 100%) and mounted in DPX.

### Alcian Blue staining

Tissue sections were rehydrated (as described above) and stained for 15 minutes in Alcian Blue solution (1% Alcian Blue in 3% solution of acetic acid) (Cat. # A5268, Sigma Aldrich, UK). Slides were then washed in running tap water for 2 minutes. Nuclei were stained in Hematoxylin solution for 1 minute and sections were then washed in tap water for two minutes. Slides were finally dehydrated (as described before), cleared and mounted in DPX.

### Elastic Van Gieson (EVG) staining

Verhoeff’s Hematoxylin solution (Cat. # RRSK40-100, Atom Scientific, UK) composed of alcoholic Hematoxylin, 10% ferric chloride and Lugol’s iodine was freshly prepared and used to stain tissue sections for 30 minutes. Slides were then washed in running tap water and differentiated with 2% ferric chloride solution, followed by a wash in distilled water. The slides were then counterstained in Van Gieson solution for 5 minutes, followed by dehydration (as described before), clearing in xylene (Cat. # 214736, Sigma Aldrich, UK) and mounting in DPX.

### Movat pentachrome (Modified Russell-Movat) staining

Tissue slides were stained using the Movat Pentachrome Stain kit (Cat. # MPS-1, Scytek, USA). Briefly, slides were left for 20 minutes in a freshly made Elastic Stain solution (5% Hematoxylin, 10 % Ferric Chloride, Lugol’s Iodine solution) and differentiated using 2% Ferric Chloride. Slides were then left for 1 minute in 5% Sodium Thiosulfate solution, before the application of Alcian blue solution pH 2.5 for 25 minutes. Biebrich Scarlet/Acid Fuchsin solution was applied for 10 minutes followed by 5% phosphotungstic acid solution and differentiation with 1% Acetic Acid solution. Finally, the slides were stained with Tartrazine solution for two minutes, dehydrated (as described before), cleared in xylene and mounted in DPX.

### Picro-sirius red staining

Tissue slides were rehydrated (as described before) and equilibrated in tap water. Samples were then stained with Picro-sirius red solution (Cat. # ab150681, Abcam, UK) for 1 hour. The slides were rinsed twice in serial baths of 0.5% Acetic Acid solutions and dehydrated in two changes of absolute ethanol. Samples were then cleared in xylene (Cat. # 214736, Sigma Aldrich, UK) and mounted in DPX. Pictures of the acellular matrix were taken under normal bright field light and birefringence mode with a planar prism.

### Lung slices culture

Acellular samples were embedded in OCT as described before, however without the fixation step in 4% PFA. The samples were cut to a thickness of 50μm with a Leica cryostat. Slices were then placed on 13mm diameter poly-lysine coated coverslips. Slices were equilibrated in PBS and then sterilised under UV light for 25 minutes. Slices were then stored at 4°C on sealed 24 well plates until needed.

### Acellular lobe isolation and sterilisation

Acellular lungs from the decellularization process were subjected to right caudal lobe isolation for further experimentation. Briefly, the left lung was occluded with 3-0 Mersilk suture silk (Cat. # W328, EVAQ8, UK), followed by the right cranial lobe, right middle lobe and accessory lobe. To assess the correct occlusion of the excluded lobes and the leaking-free inflation of the right caudal lobe, the airways and vasculature were perfused with 1ml of PBS. The isolated lobes were placed in PBS P/S solution to be sterilised by γ irradiation.

### Cell seeding

A549 or HBEC cells were cultured in their respective expansion medium prior seeding on lung scaffold. HBECs were a kind donation from Samuel Janes’ lab (UCL, London). HBECs were cultured on mitotically inactivated 3T3-J2 feeder cells in HBEC maintenance medium (described below). Epithelial cells were cultured at 37°C and 5% CO_2_ with two to three medium changes a week. To isolate only the HBEC population, a differential dissociation was performed, taking advantage of the greater sensitivity of feeder cells to trypsin in comparison with strongly adherent epithelial cells. All dissociations were performed with 0.05% Trypsin-EDTA (Cat. # 25300, Gibco, UK). When optimal confluency was reached, epithelial cells were differentially detached, counted and passed through a 40μm cell strainer to ensure a single cell suspension. A minimum of 20 million A549 or HBEC were then manually seeded through the airways in a volume of 800μl epithelial maintenance medium using a 1ml syringe. In the case of dynamic culture conditions, mechanical medium ventilation was initiated from overnight to 18 hours after seeding. HBEC maintenance medium was changed every other day, and cells were cultured for 4 days, both under static and dynamic culture conditions.

### HBEC maintenance medium

HBEC culture medium consisted of DMEM (Cat. # 41966, Gibco, UK) and F12 (Cat. # 21765, Gibco, UK) (3:1 ratio), penicillin–streptomycin (P/S) (Cat. # 15070, Gibco, UK) and 5% FBS (Cat. # Gibco, UK) supplemented with 5mM Y-27632 (Cat. # Y1000, Cambridge Bioscience, Cambridge, UK), 25ng/ml hydrocortisone (Cat. # H0888, Sigma Aldrich, St. Louis, MO), 0.125ng/ml epidermal growth factor (EGF) (Cat. # 10605, Sino Biological, Beijing, China), 5mg/ml insulin (Cat. # I6634, Sigma Aldrich, UK), 0.1nM cholera toxin (Cat. # C8052, Sigma Aldrich, UK), 250ng/ml amphotericin B (Cat. # 10746254, Fisher Scientific, Loughborough, UK) and 10μg/ml gentamicin (Cat. # 15710, Gibco, UK).

### A549 maintenance medium

A549 culture medium consisted of DMEM (Cat. # 41966, Gibco, UK), 10% Foetal Bovine Serum (FBS) (Cat. # 10270106, Gibco, UK), 2mM L-Glutamine (Cat. # 25030081, Thermo Fisher, UK) and 100U/ml Penicillin-Streptomycin (Cat. # 10378016, Thermo Fisher, UK).

### Viral infection

The bioengineered lungs were infected with Respiratory Syncytial Virus (RSV), the one used for this study is GFP labelled, to facilitate the detection of infected cells. Recombinant GFP tagged RSV A2 strain was kindly provided by Fix et al. (10.2174/1874357901105010103). Each lung was exposed to a RSV at a multiplicity of infection (MOI) of 0.1. MOI was calculated by PFU/ml of RSV virus in reduced Serum Medium (Opti-MEM 31985062) per number of cells seeded per lung. The inoculum was added via tracheal injection with a 1ml Leur-lok syringe. After the infected lung was incubated for 1 h at 37°C in static condition, the bioreactor was restarted to proceed with 4 days of dynamic culture. Plastic static controls of infection were maintained at the same condition as the bioreactor to monitor the state of infection.

### Drug testing

After the infection the lungs were treated with Remdesivir(Bio-Techne Catalog # 7226), an antiviral nucleotide analogue. The drug stock was prepared as 10 mg/ml in DMSO and then diluted to a final concentration of 8 μg/ml in culture media for each treated bioreactor. Control lungs were mock infected with the same volume of DMSO alone in culture media.

### Quantification of RSV by RT-PCR

RNA was extracted from media samples using QIAamp Viral RNA Mini Kit (250) (Qiagen) as per the manufacturer’s instructions. RNA was quantified using the Nanodrop 1000 (ThermoFisher). cDNA was Synthesised using the High-Capacity RNA-to-cDNA kit (Applied Biosystems) as per the manufacturer’s instruction and using a starting concentration of 0.5μg total RNA in a final reaction volume of 20μl. A no reverse transcriptase control was also performed to control for remaining contaminating DNA in the RNA preparation. Quantitative RT-PCR was performed using TaqMan Universal Master Mix II, with UNG (Applied Biosystems). A total reaction volume of 20μl with 6μl of cDNA per reaction was used. Primers and the probes were designed and used as previously described[29][30] (listed in Supplementary Methods). Primers were used at a final concentration of 10mM and probes used at a final concentration of 2pM. Samples were run on an AB Biosystems step one plus RT-PCR machine. Reaction conditions-1 cycle 50°C for 2 minutes, 1 cycle 95°C for 10 minutes, 40 cycles 95°C for 15 seconds followed by 60°C for 1 minute. A template plasmid containing the N protein sequence[31] was used to quantify the N protein copy number in media samples. RSV load was extrapolated from a standard curve of known N protein copies. The number of RSV genome copies in infected media samples was extrapolated from the standard curve.

### Quantification of infectious RSV particles by plaque assay

The day prior to infection 1.15-1.5 x 105 Vero E6 cells were seeded in 0.5 ml Media (DMEM with 10% FCS,1% Pen/Strep) per well of 24 well plate. On the day of infection, media samples were serially diluted as 10-fold dilutions in OptiMEM. Media from each well was replaced with 200ul of sample. Control wells received viral inoculum containing 200 PFU/ml or mock infected media samples alone. Vero E6 cells were incubated with inoculum in a 37oC CO2 incubator for 1 hour. Inoculum was then aspirated and replaced with 0.5 ml of 1:1 2.4% microcrystalline cellulose suspension: 2x MEM (+4% FCS, LG, Pen/Strep, pH 7.35). After 7 days in a 37oC CO2 incubator, cells were fixed with 0.2ml of 2% PFA + 2% glutaraldehyde (PolySciences) per well and subsequently stained with 0.2ml of crystal violet solution (8% crystal violet + 20% ethanol in ddH2O). Plaques were counted and averaged between technical repeats. Each plaque represents one plaque forming unit (PFU), this was calculated per ml by multiplying the plaques/well by dilution factor and given per total in circulating bioreactor media of 80ml (PFU/bioreactor).

## Notes

### Competing Interest Statement

The authors have declared no competing interest.

