## Supplementary material for "Lung viral infection modelling in a bioengineered whole-organ": Suppplementary informations

### Supplementary tables

| Primary antibody/ Epitope | Host species | Dilution | Supplier (Cat. #) |
|---|---|---|---|
| Ki67 | Rabbit | 1:400 | Abcam (ab15580) |
| Caspase 3 | Rabbit | 1:300 | Cell Signalling (9661S) |
| FOXP2 | Rabbit | 1:200 | Abcam (ab16046) |
| NKX2.1 | Rabbit | 1:200 | Abcam (ab76013) |
| Laminin | Rabbit | 1.200 | Abcam (ab11575) |
| E-Cadherin | Rabbit | 1:100 | BD (BD610181) |
| Keratin 8 | Rabbit | 1:100 | Abcam (ab59400) |
| Keratin 18 | Mouse | 1:100 | Abcam (ab668) |
| Keratin 5 | Mouse | 1:100 | Abcam (ab128190) |
| Keratin 14 | Rabbit | 1:800 | Biolegend (PRB-155P) |
| Integrin α6 | Mouse | 1:200 | Abcam (ab20142) |
| p63 | Rabbit | 1:200 | Abcam (ab124762) |
| Collagen I | Mouse | 1:200 | Abcam (ab6308) |
| Collagen IV | Rabbit | 1:200 | Abcam (ab6586) |

| Secondary antibody | Host species | Dilution | Supplier (Cat. #) |
|---|---|---|---|
| αMouse Alexa Fluor 488 | Goat | 1:200 | Thermo Fisher (A-11001) |
| αRabbit Alexa Fluor 594 | Goat | 1:200 | Thermo Fisher (A-11012) |

| Gene | Primer sequence |
|---|---|
| CK14 | Forward: CATGAGTGTGGAAGCCGACAT |
| | Reverse: GCCTCTCAGGGCATTCATCTC |
| CK5 | Forward: GGA GTT GGA CCA GTC AAC ATC |
| | Reverse: TGG AGT AGT AGC TTC CAC TGC |
| ΔN p63 | Forward: AAAGGACAGCAGCATTGATCAA |
| | Reverse: TGTTCAGGAGCCCCAGGTT |

| Gene | Primer sequence 5’-3’ |
|---|---|
| RSV F | CTCAATTTCCTCACTTCTCCAGTGT |
| RSV R | CTTGATTCCTCGGTGTACCTCTGT |
| RSV PROBE | TCCCATTATGCCTAGGCCAGCAGCA |

### Supplementary Figure 1

A

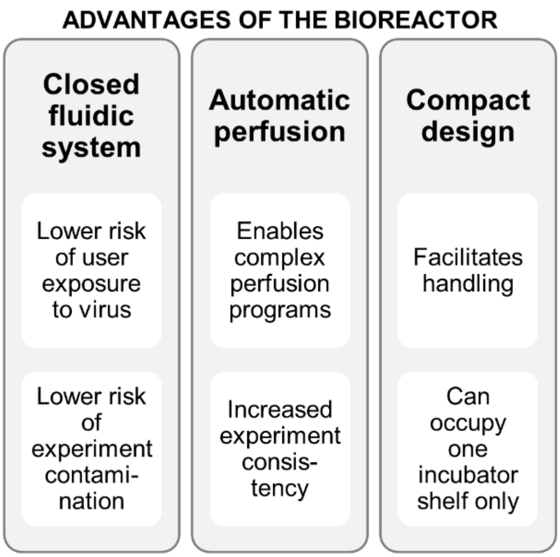


B

| | DECELLULARISATION FLOW RATES [ml/min] | | |
|---|---|---|---|
| | Lung IN (P1) | Lung OUT (P2) | Priming/Extraction |
| n1 | 11.00 | 5.06 | 13.40 |
| n2 | 11.20 | 5.10 | 13.20 |
| n3 | 11.00 | 5.20 | 13.20 |
| **μ** | **11.07** | **5.12** | **13.27** |
| **σ** | **0.115** | **0.072** | **0.115** |

| | RECELLULARISATION FLOW RATES [ml/min] | | |
|---|---|---|---|
| | Medium Perfusion | Level Control | Priming |
| n1 | 2.48 | 10.50 | 5.50 |
| n2 | 2.46 | 10.50 | 5.30 |
| n3 | 2.52 | 10.40 | 5.40 |
| **μ** | **2.49** | **10.47** | **5.40** |
| **σ** | **0.031** | **0.058** | **0.100** |

C / D

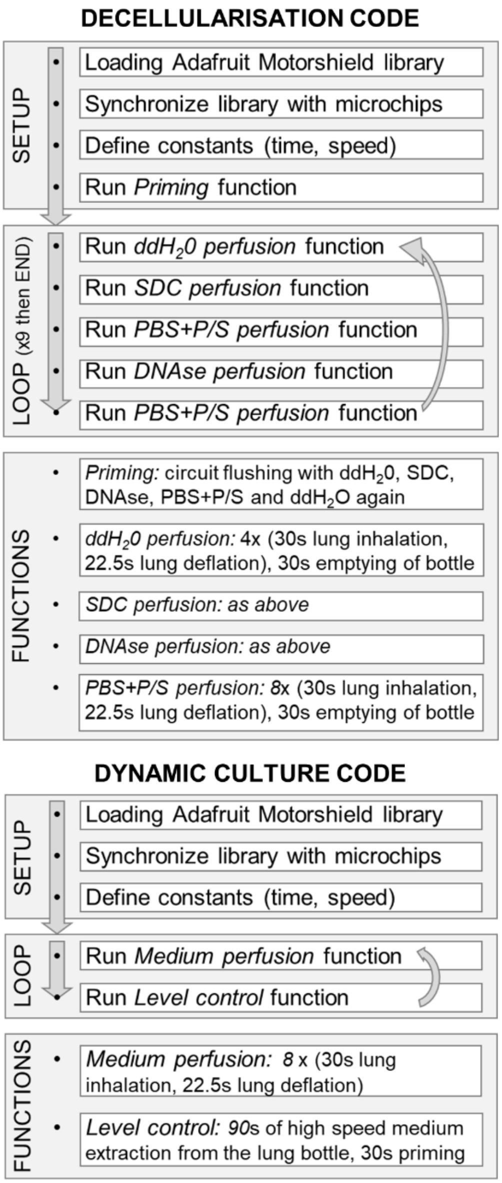


E

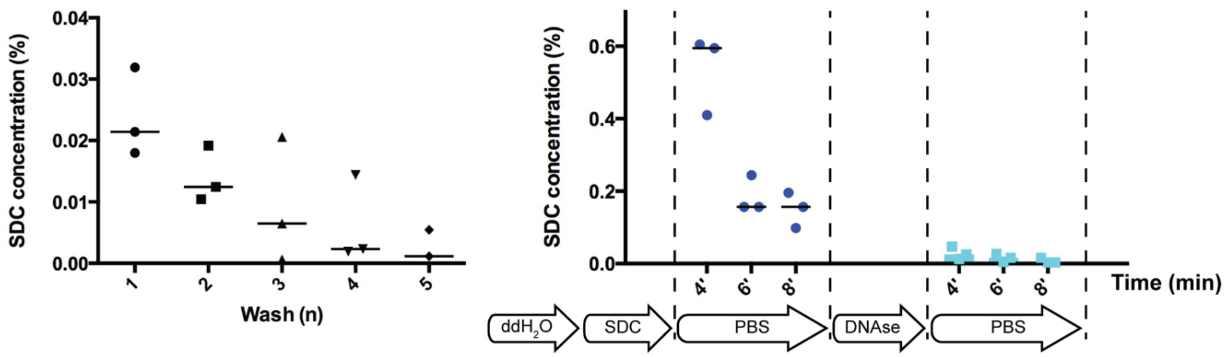

**Supplementary Figure 1. A.** Table summarising the main operational advantages of the bioreactor system. **B.** Table showing the flow rates measured during decellularization and dynamic culture programs. **C.** Arduino code box diagram showing the main steps of the decellularization program, namely priming and cyclic perfusion. **D.** Arduino code box diagram showing the main steps of the recellularization process, namely medium perfusion, level control and priming. **E.** Graphs showing the amount of residual SDC within decellularized scaffolds, measured by its level within PBS lung washes during the decellularization process.

### Supplementary Figure 2

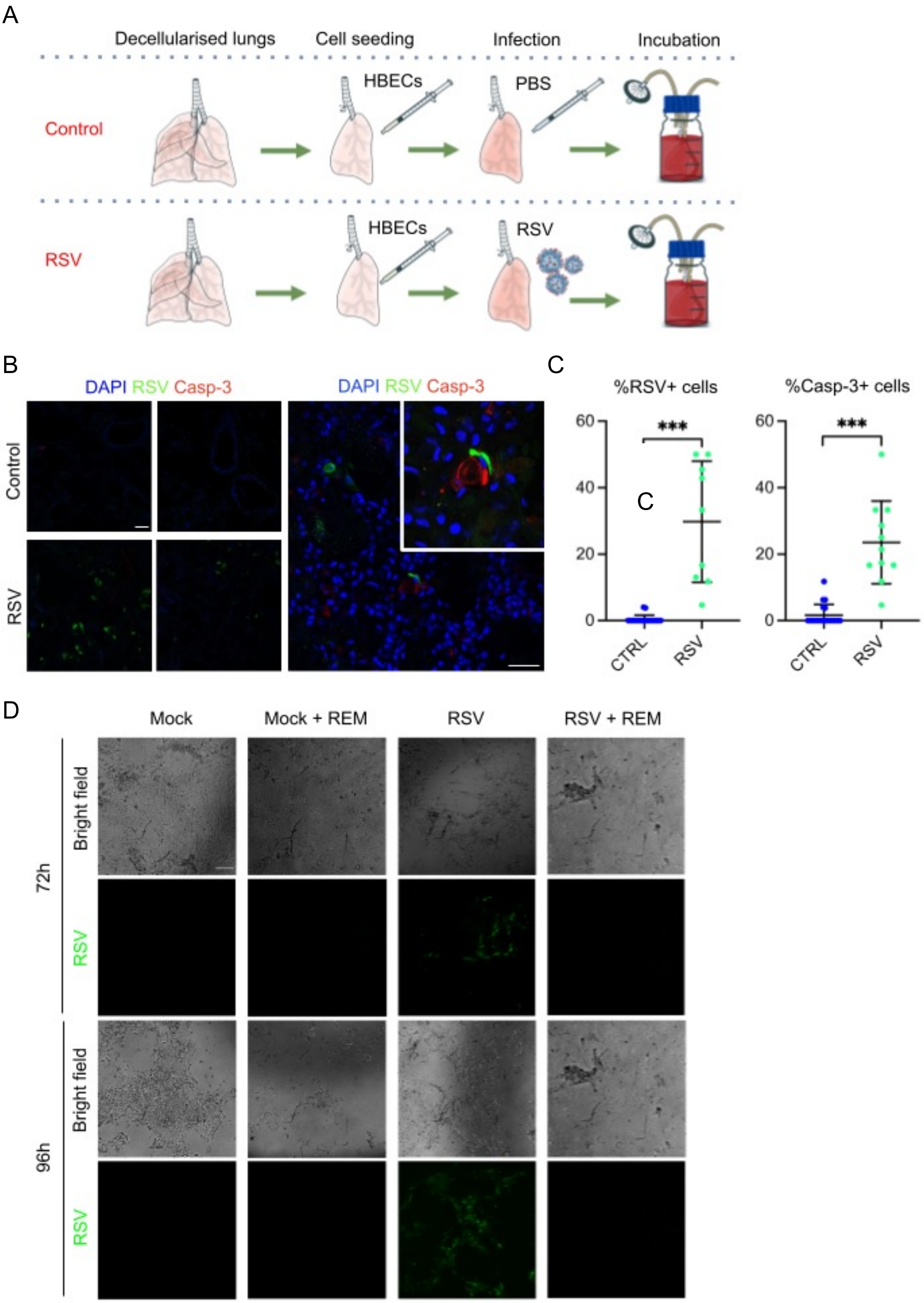

**Supplementary Figure 2. A**. Preliminary experimental setup for the recellularization and infection of the engineered lung with an RSV virus. **B**. Immunofluorescence analysis of nuclei, RSV, Caspase-3 in the engineered lungs cultured under the two experimental conditions. Scale bar 100µm. **C.** Quantification of the immunofluorescence analysis in C confirms a significant increase in the percentage of RSV-positive cells in the RSV infection condition (left), along with an increase in the percentage of Casp3-positive cells. T-test; ***p<0.005. **D.** Bright field and fluorescence images of cells cultured in Petri Dish as control of the experiment described in main Figure 4B after 72 and 96 hours post-infection. Scale bar 100µm.
